# Inactivating Negative Regulators of Cortical Branched Actin Enhances Persistence of Single Cell Migration

**DOI:** 10.1101/2023.05.13.540631

**Authors:** Artem I. Fokin, Arthur Boutillon, John James, Laura Courtois, Sophie Vacher, Gleb Simanov, Yanan Wang, Anna Polesskaya, Ivan Bièche, Nicolas B. David, Alexis M. Gautreau

## Abstract

The Rac1-WAVE-Arp2/3 pathway pushes the plasma membrane by polymerizing branched actin at the cell cortex and thereby powering membrane protrusions that mediate cell migration. Here, using knock-down (KD) or knock-out (KO), we combine the inactivation of the Arp2/3 inhibitory protein Arpin, the Arp2/3 subunit ARPC1A and the WAVE complex subunit, CYFIP2, that all enhance the polymerization of cortical branched actin (CBA). Inactivation of the 3 CBA negative regulators increases migration persistence of human breast MCF10A cells, and of endodermal cells in the zebrafish embryo, significantly more than any single or double inactivation. In the triple KO, but not triple KD cells, the “super-migrator” phenotype was associated with a heterogenous down-regulation of vimentin expression and a lack of coordination in collective behaviors, such as wound healing and acinus morphogenesis. Re-expression of vimentin in triple KO cells restored the normal persistence of single cell migration to a large extent, suggesting that vimentin down-regulation is one of the adjustments in gene expression through which the super-migrator phenotype is stably maintained in triple KO cells. Constant excessive production of branched actin at the cell cortex thus commits cells into a motile state through changes in gene expression.

## INTRODUCTION

Cell migration is a critical physiological process during embryo morphogenesis, especially during gastrulation, and in some cell types of the adult, such as immune cells patrolling the organism. Most adult cells, however, do not migrate. They, nonetheless, can be induced to migrate in pathological conditions, for example during cancer progression. Untransformed cells classically use the mesenchymal mode of cell migration that relies on the formation of adherent membrane protrusions called lamellipodia. Lamellipodia are powered by polymerization of branched actin by the Arp2/3 complex (Ridley, 2011; Wu et al., 2012). Arp2/3 activity is regulated by Nucleation-Promoting Factors (NPFs), the most important of which for membrane protrusions and cell migration being WAVE (Bieling and Rottner, 2023). WAVE is embedded into a multiprotein complex, which is activated by the small GTPase Rac1 at the leading edge of cells (Ding et al., 2022; Rottner et al., 2021). Branched actin is thought to feedback on Rac1 activation and thus to sustain polymerization of branched actin where it was previously polymerized (Castro-Castro et al., 2011; Krause and Gautreau, 2014). Thereby the Rac1-WAVE-Arp2/3 pathway controls persistence of cell migration.

Many proteins inhibit the Rac1-WAVE-Arp2/3 pathway at different levels. Arp2/3 activation can be blocked by several Arp2/3 inhibitory proteins (Chánez-Paredes et al., 2019; Zhao et al., 2020). Arpin occupies one of the two NPF binding sites of Arp2/3 and thus prevents WAVE from activating it (Fregoso et al., 2022). Therefore, Arpin depletion by KD or KO promotes migration persistence (Dang et al., 2013; Simanov et al., 2021). Branched actin networks can also be debranched by coronins and GMF family proteins that recognize Arp2/3 at the branched junction (Molinie and Gautreau, 2018). The WAVE regulatory complex is inhibited by the Nance-Horan Syndrome family protein NHSL1 that interacts with the WAVE complex and can even replace WAVE in its complex (Law et al., 2021; Wang et al., 2022). Specific GTPase activating proteins inhibit the small GTPase Rac1 to restrict lamellipodial protrusions and cell migration (Parrini et al., 2011; Yamazaki et al., 2013). The ability of GTP-bound Rac1 to activate the WAVE regulatory complex is restricted by CYRI proteins that compete with WAVE complexes for Rac1 binding (Fort et al., 2018; Yelland et al., 2021).

The combinatorial complexity in the assembly of WAVE and Arp2/3 complexes provides additional means to regulate membrane protrusions and cell migration. The paralogous subunits ARPC1B and ARPC5L assemble Arp2/3 complexes that are more active than the ones assembled with ARPC1A and ARPC5 (Abella et al., 2016). We have previously shown that depletion of ARPC1A allows the assembly of more ARPC1B-containing complexes, because common subunits are no longer distributed between the two paralogous proteins, and that ARPC1B-containing Arp2/3 complexes promote cortical branched actin and migration persistence (Molinie et al., 2019). Similarly, we have shown that the CYFIP2-containing WAVE complexes are less readily activated by Rac1 than CYFIP1-containing WAVE complexes and that CYFIP2-depleted cells display increased lamellipodial protrusions and migration persistence (Polesskaya et al., 2022). Because of this balance of paralogous subunits, ARPC1A and CYFIP2 exhibit an apparent inhibitory activity on membrane protrusion and migration persistence, even though these subunits belong to complexes that promote branched actin polymerization. Cortical branched actin polymerized by the Rac1-WAVE-Arp2/3 pathway does not only control cell migration, but also the decision to enter a new cell cycle. In single cells, migration persistence was found to inversely correlate with the duration of the G1 phase (Molinie et al., 2019).

In epithelial cells, cell-cell junctions also depend on the Rac1-WAVE-Arp2/3 pathway for their assembly and maintenance (Li et al., 2020; Verma et al., 2012). When cells reach a high density, they suppress migration and proliferation (Puliafito et al., 2012; Streichan et al., 2014). Upon wounding the monolayer, cells resume migration and proliferation to heal the wound in a coordinated manner regulated in time and space (Palamidessi et al., 2019; Poujade et al., 2007). Because of this coordination between cells, wound healing is a more complex process than single cell migration. Epithelial cells can also detach from each other through an Epithelial to Mesenchymal Transition (EMT) that relies on changes of gene expression (Thiery et al., 2009). It appeared that there are multiple partial EMT states that determine migration modes (Nieto et al., 2016). Vimentin is an intermediate filament protein that is a marker of EMT. It is initially widely expressed in the embryo and becomes restricted to mesenchymal cells (Paulin et al., 2022). Vimentin is involved in cell migration and appears critical for wound healing in vivo (Eckes et al., 1998; Eckes et al., 2000).

An important question is thus how negative regulators of WAVE-dependent polymerization of branched actin maintain the migration of epithelial cells under control. Here we show that enhancing polymerization of branched actin at the cell cortex by removing as many as 3 negative regulators promotes single cell migration, but not collective migration of mammary epithelial cells. Cells adapt to these long-term perturbations by altering gene expression, in particular, by down-regulating vimentin expression. Vimentin down-regulation enhances migration persistence of single epithelial cells.

## RESULTS

### Knocking-out 3 CBA negative regulators greatly increases migration persistence

We recently characterized the role of the RAC1-WAVE-Arp2/3 pathway in sustaining migration in a direction chosen at random by single MCF10A cells (Molinie et al., 2019; Polesskaya et al., 2022). The human MCF10A cell line is derived from a fibrosis of mammary breast (Soule et al., 1990). Cells are diploid and not transformed, since they do not form tumors when grafted into immunocompromised mice (Worsham et al., 2005). These epithelial cells are quite plastic, as single cells detach from epithelial islets in vitro and adhere again to other cells when they meet. Arpin, ARPC1A and CYFIP2 play negative roles towards the polymerization of cortical branched actin (CBA) at different levels of the RAC1-WAVE-Arp2/3 pathway (Fig. 1A) and were inactivated.

**Figure 1.**
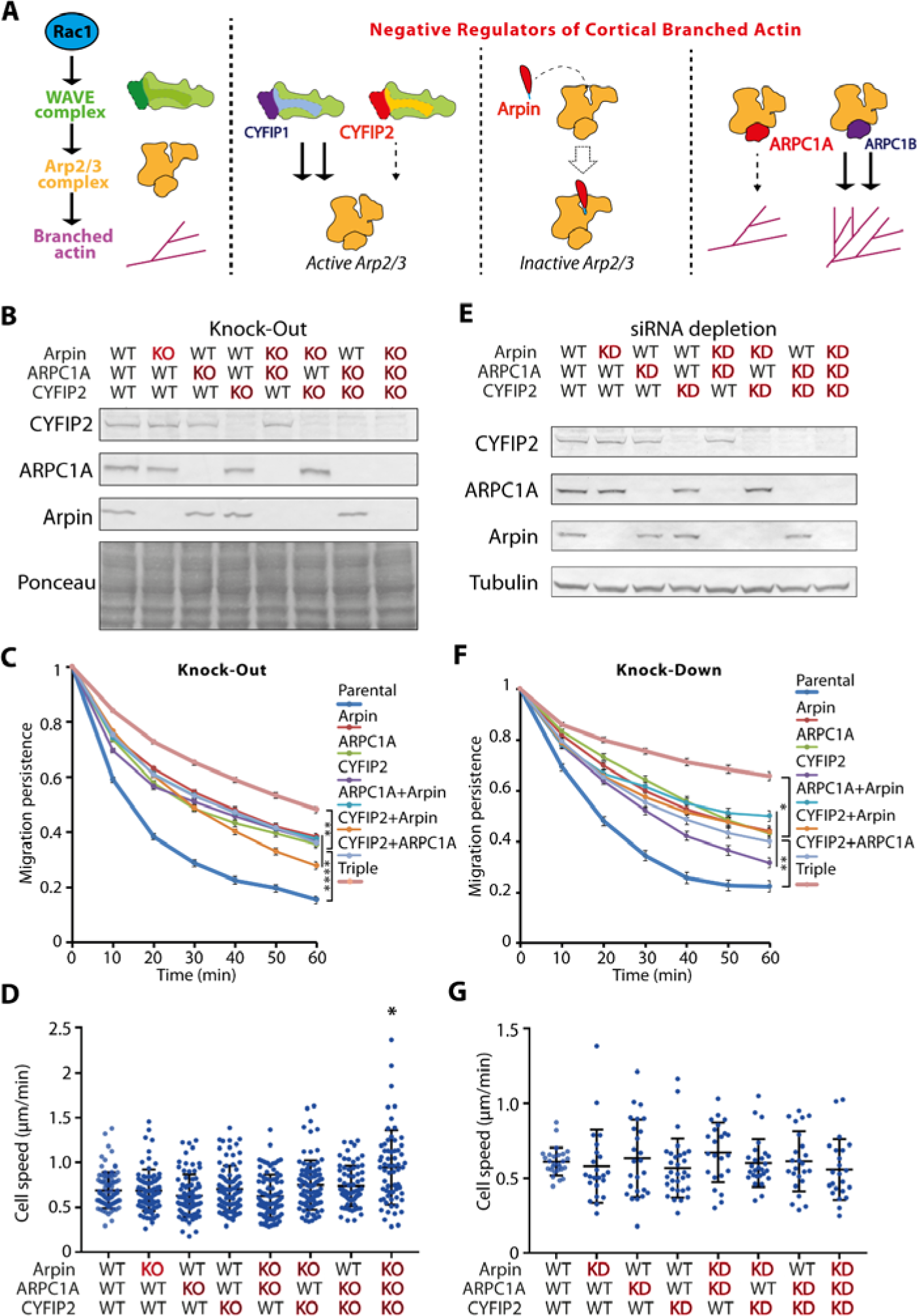
Simultaneous KO or KD of 3 CBA negative regulators increases migration persistence. **A** Scheme representing the role of each CBA negative regulator, Arpin, ARPC1A and CYFIP2 in the Rac1-WAVE-Arp2/3 pathway. **B** MCF10A cells were depleted of the 3 CBA negative regulators in various combinations using Cas9-mediated knock-out (KO). Western blots of target proteins. **C** Migration persistence of KO cells. **D** Average speed of KO cells (mean ± sem). **E** MCF10A cells were depleted of the 3 CBA negative regulators in various combinations using siRNAs. Western blots. **F** Migration persistence of knock-down (KD) cells **G** Average speed of KD cells (mean ± sem). Mean ± sem, 3 independent experiments for both KD and KO, n > 21 for KD, n > 63 for KO. ANOVA, * p<0.05, ** p<0.01, **** p<0.0001.

To generate KO clones for each of these CBA negative regulators, we have created insertions/deletions (indels) in the Open Reading Frame (ORF) by transfecting synthetic gRNAs with the purified Cas9 protein into MCF10A cell line. In this protocol with no antibiotic selection, screening of about hundred clones by Western blot allowed to isolate at least two independent KO clones for each of these genes (Molinie et al., 2019; Polesskaya et al., 2022). We analyzed genomic DNA of these clones to characterize the indels. Genomic DNA encompassing the Cas9-mediated cut was amplified and the PCR product was sequenced. The PCR sequence could be directly read if the two alleles were the same. This was the case of ARPIN KO clone #1. If the PCR sequence cannot be read, it indicates that the two alleles generate different sequences that overlap. In this case, the PCR product was cloned and individual plasmids were then sequenced to unambiguously determine the sequence of the two alleles. In all clones, we were able to identify the indels that accounted for the 2 KO alleles due to frameshift (Fig.S1). For each of these CBA negative regulators, the two KO clones had increased migration persistence, although to a different extent (Fig.S2). This systematic increase of migration persistence was not associated with consistent variations of cell speed or Mean Square Displacement (MSD), as we previously reported for MCF10A cells when the RAC1-WAVE-Arp2/3 pathway was perturbed (Molinie et al., 2019; Polesskaya et al., 2022).

To explore the potential synergistic role of regulatory proteins, we combined their KOs. Our goal was to see whether we could isolate a “super-migrator” cell line, since our protocol of gene inactivation, which does not require antibiotic selection, allows to perform the whole procedure again in a previously obtained KO cell line. For each CBA negative regulator, we edited further the KO clone that displayed the most increased migration persistence (Fig.S3). We managed to obtain the 3 possible double KOs, ARPIN ARPC1A, ARPIN CYFIP2 and ARPC1A CYFIP2 (Fig.1B). To our surprise, none of the double KOs migrated more persistently than single KOs (Fig.1CD, Fig.S3). We thus attempted to combine the 3 KOs and this time, we obtained a super-migrator cell line that migrated more persistently than any single or double KO of ARPIN, CYFIP2, and ARPC1A (Fig.1CD, Fig.S3, Movie S1).

A difficulty associated with this approach is that we could not afford to systematically study two independent clones for combined KOs, even though we had observed significant differences between single KO clones (Fig.S2). This rule of keeping two independent clones for each genotype would mean 4 clones for each of the 3 double KOs and 8 clones for the triple KO. Moreover, if a clone adapts to its genotype, then the route taken to sequentially introduce mutations might potentially also affect the phenotype. In other words, the sequence by which compound KO were obtained might also influence the phenotype. To test all possible routes would double the number of double KO clones and multiply by 6 the number of triple KO clones. To be fully rigorous, the total number of clones to compare with parental cells would amount to 4×2 + 4×2 + 8×6 = 64 clones. This large number of clones prompted us to compare the super-migrator cell line we had obtained to the transient depletion of the 3 CBA negative regulators.

### Knocking-down the 3 CBA negative regulators reveals a super-migrator phenotype in cell culture and zebrafish embryos

We transiently transfected MCF10A cells with pools of siRNAs targeting each of the 3 CBA negative regulators (Fig.1E). Upon siRNA-mediated depletion, double Knock-Down (KD) cells had the same increased migration persistence as single KD cells (Fig.1F). Only the triple KD had increased persistence compared with all single or double KDs. The phenotype of the triple KD is thus most similar to the triple KO clone we isolated. A difference between the two approaches is that triple KO cells had a slight increase of cell speed, which was not exhibited in triple KD cells (Fig.1G). Together these experiments showed that MCF10A cells greatly increased migration persistence upon down-regulation of 3, but not 2 CBA negative regulators, whether this down-regulation was performed by KO or KD.

To evaluate the physiological relevance of these observations, we turned to zebrafish embryos. We have previously characterized the migration of endodermal cells, which internalize at the beginning of gastrulation and then disperse over the yolk surface as single cells through random walks (Pézeron et al., 2008). This cell-autonomous migration process is governed by the Rac1 – Arp2/3 pathway (Giger and David, 2017; Woo et al., 2012). Using morpholinos, we knocked down Arpin, ARPC1A and CYFIP2 in endodermal cells and transplanted some into receiver zebrafish embryos (Fig.2A). Transplanted cells were tracked thanks to the expression of a Histone2B-mCherry fusion (Fig.2B, Fig.S5, Movie S2).

**Figure 2.**
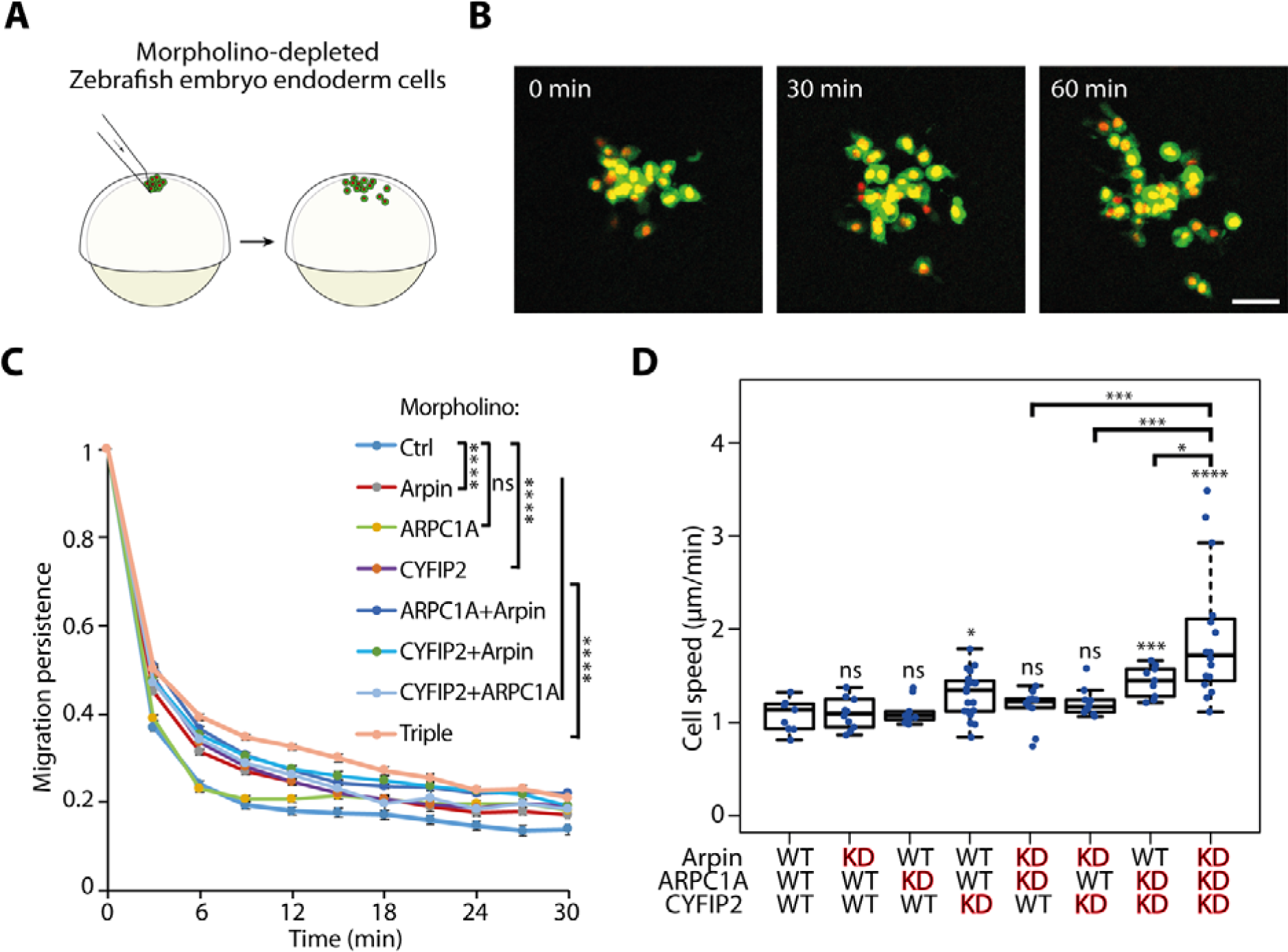
Enhanced migration of endodermal cells in Zebrafish embryos upon morpholino-mediated KD of CBA negative regulators. **A** Transplantation of Histone 2B-mCherry-labeled endodermal cells from a gsc:GFP transgenic donor embryo to a receiver embryo allows to monitor migration of single endodermal cells. **B** Dissemination of labeled cells over time. Scale bar 50 *μ*m. **C** Migration persistence of endodermal cells. **D** Average speed of endodermal cells (mean ± sem). Number of embryos/average number of cells in each embryo per condition: Ctrl 9/58, Arpin 9/32, ARPC1A 10/32, CYFIP2 20/39, ARPC1A+Arpin 10/23, CYFIP2+Arpin 9/58, CYFIP2 + ARPC1A 10/13, Triple 16/14. An average of 33 cells were transplanted in each embryo. ANOVA, * p<0.05, ** p<0.01, *** p<0.001, **** p<0.0001, n.s. non-significant. When not indicated otherwise, p refers to the comparison with the WT condition.

When CBA negative regulators were tested in isolation, KD of CYFIP2 or Arpin significantly increased migration persistence of endodermal cells (Fig.2C), as we previously reported in the collective migration of prechordal plate cells (Dang et al., 2013; Polesskaya et al., 2022). Double KDs of Arpin and CYFIP2 led to an even stronger increase in persistence. ARPC1A KD had no effect on the migration of endodermal cells. Consistently, KDs of ARPC1A and CYFIP2 together led to a phenotype similar to simple CYFIP2 KD. But KDs of ARPC1A and Arpin increased cell persistence compared with the single KD of Arpin, as if ARPC1A potentiated the effect of Arpin. The triple depletion of Arpin, ARPC1A and CYFIP2 rendered cells significantly more persistent than all single or double depletions (Fig.2C, Fig.S5). Triple KD cells were also faster than other cells (Fig.2D). In conclusion, results observed in the zebrafish embryo largely mirror those obtained on MCF10A cells in culture, suggesting that KDs of the 3 CBA negative regulators turn cells into super-migrators in vivo as well as in culture cells.

### The population of super-migrator triple KO cells is heterogeneous

We then asked whether turning on migration persistence would be associated with defects. We thus decided to challenge our MCF10A KO clones into a morphogenetic assay, where single mammary cells proliferate and develop acini at the surface of matrigel (Debnath et al., 2003). We have previously reported that Arpin KO formed normal hollow acini, albeit bigger than parental cells due to increased proliferation (Molinie et al., 2019). This was also the case upon KO of ARPC1A or CYFIP2 (Fig.3A-B). Acinus sizes reached by double KOs were not different from the ones of single KOs, but the biggest acini from triple KO reached a size bigger than the ones differentiated from single and double KOs. Parametric statistics could not be applied, because the size distribution of acini differentiated by the triple KO became scattered, with coexistence of small and large acini. Most large acini from the triple KO developed a lumen, more frequently so than the acini differentiated from parental cells (Fig.3C-D). Most small acini of the triple KO did not develop a lumen, in line with a delayed morphogenesis. Some of these small acini, however, displayed an irregular shape together with heterogeneous laminin deposition (Fig.3C). In conclusion, excessive activation of the Rac1-WAVE-Arp2/3 pathway did not only increase cell proliferation, as previously reported (Molinie et al., 2019), but also increased the heterogeneity of the cell population.

**Figure 3.**
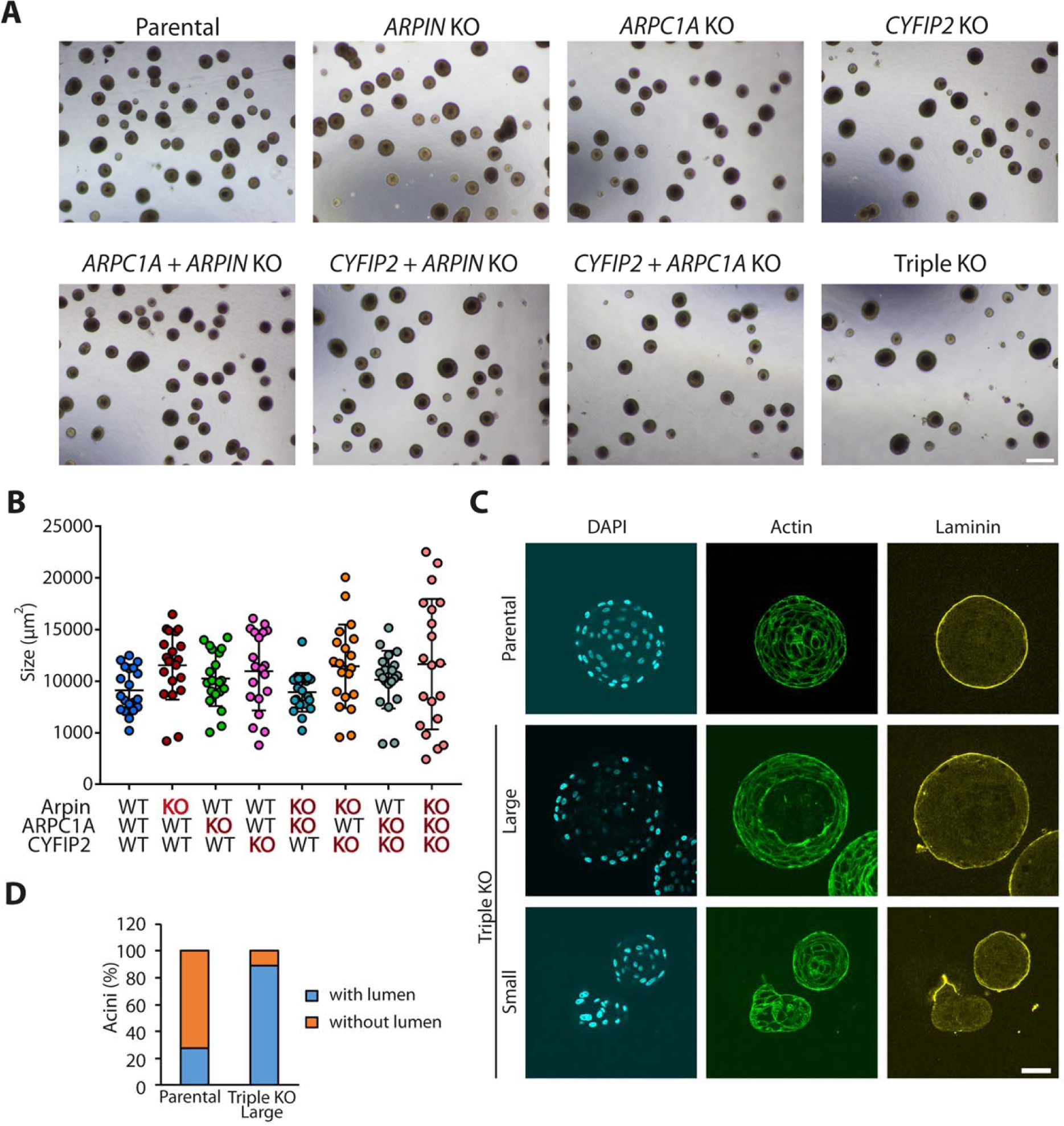
Inactivation of CBA negative regulators alter acini morphogenesis. **A** Overview of MCF10A acini growing on top of matrigel. Scale bar 200 μm. **B** Acini size. Apparent area (mean ± sd) is plotted. Bigger acini were formed when more CBA inhibitors were inactivated, but size heterogeneity also increases. **C** MCF10A acini derived from parental and triple KO cells were stained with DAPI and phalloidin to visualize nuclei and actin filaments and with laminin antibodies to reveal the extracellular matrix secreted by MCF10A acini. In Triple KO, large acini often contained a single lumen, whereas small acini had no lumen. Small acini tend to deposit laminin irregularly and display irregular shapes. Scale bar 50 μm. **D** Quantification. n = 26 acini for parental and n = 22 acini for Triple KO.

We then sought to analyze the triple KO for collective cell migration in a wound healing assay to examine how cells would coordinate with each other during cell migration, since Rac1-dependent polymerization of branched actin is critical both for lamellipodial protrusions at the front edge and for the maintenance of adherens junctions (Fenteany et al., 2000; Verma et al., 2004; Verma et al., 2012). The triple KO line that was super-migrating at the single cell level, did not improve wound healing, and even slightly decreased it, if anything (Fig.4A-B). The migrating front of the triple KO, however, appeared different from the one of parental cells (Movie S3). MCF10A cells closed the wound with a front that homogeneously progresses, unlike MDCK cells that organize multicellular ‘fingers’ pulled by leader cells (Poujade et al., 2007). The triple KO in MCF10A cells appeared more similar to MDCK cells than parental MCF10A cells, with protruding fingers at the front. To quantify this effect, we measured the distance covered by the front over time. Overall, the mean value was quite similar between parental and triple KO cells, however, the variance was higher in triple KO than in parental cells due to these fingers at the leading front (Fig.4C).

**Figure 4.**
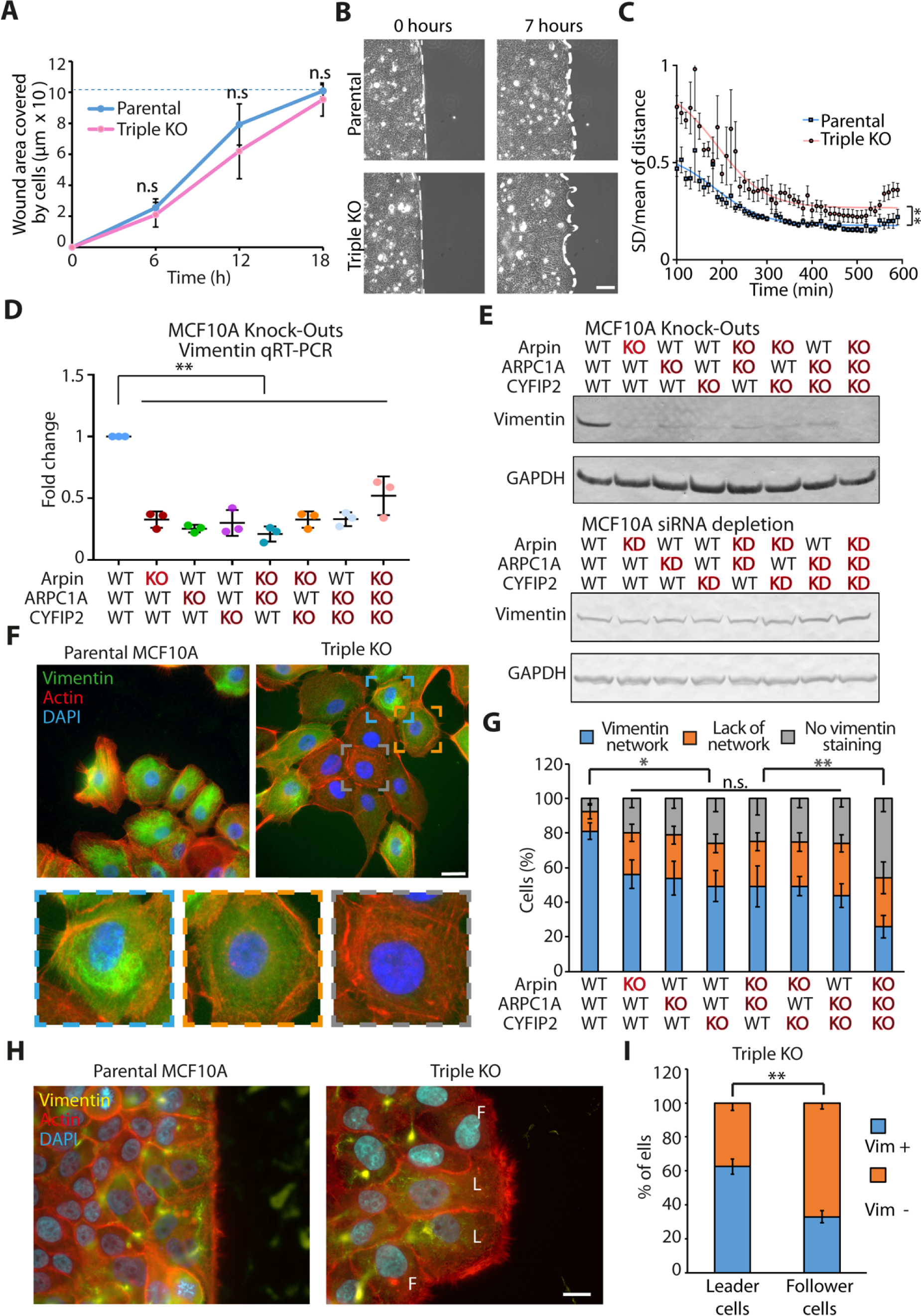
Altered collective behaviors and heterogeneity of vimentin expression in MCF10A cells inactivated for CBA negative regulators. **A** Wound healing in triple KO and parental cells. Mean ± SD of 18 wounds (3 fields of view per condition from 6 independent experiments). Kruskal-Wallis test. **B** Heterogeneous behaviors of triple KO cells at the migration front. Fingers develop at the edge of triple KO monolayers, but not of parental cell monolayers. Scale bar 100 μm. **C** Quantification of edge fluctuations. The position of front cells is reported over time and compared to the initial position to calculate the distance traveled by front cells. The ratio of SD over mean of this value is calculated for multiple fields of view (n = 6) and plotted against time after wounding. Mann-Whitney test over areas under curve. **D** Relative levels of vimentin mRNA in KO cell lines. qPCR was performed in triplicates and normalized by the mRNA level of GAPDH. Fold change in each replicate is shown as individual data points and mean ± sd. ANOVA, followed by the Dunnett’s multiple comparisons test. **E** Vimentin is down-regulated upon KO of CBA negative regulators, but not KD. Western blots. **F** Parental and triple KO cells were stained with DAPI, phalloidin (actin), and vimentin antibodies. The population of triple KO cells is heterogeneous in vimentin organization. 3 categories of cells are distinguished in the triple KO population and illustrated by a magnified representative example. Scale bar 20 μm. **G** Quantification of the 3 vimentin categories. For each condition, more than 400 cells in 11 fields of view were pooled from 2 independent experiments and analyzed. Kruskal-Wallis test. **H** Parental and triple KO cells were stained with DAPI, phalloidin (actin), and vimentin antibodies, 7 h after wounding. In triple KO monolayers, leading cells (L) in finger-like protrusions are distinguished from neighboring followers cells (F). **I** Vimentin content of leader and follower cells of triple KO. Quantification of 109 L and F cells from 3 independent wounds. Welch’s unequal variance t-test. n.s. non-significant, * p<0.05, ** p<0.01.

We then decided to examine whether KO of CBA negative regulators would affect gene expression. We first analyzed all the genes of the Rac1-WAVE-Arp2/3 pathway in all KO cell lines that we had isolated for potential compensatory changes in gene expression. We found that, in line with non-sense mRNA mediated decay due to premature stop codons (Popp and Maquat, 2016), the levels of *CYFIP2, ARPIN* and *ARPC1A* mRNAs were greatly decreased when they contained indels in KO cell lines (Fig.S6). Down-regulation of CYFIP2 and ARPC1A was not compensated by transcriptional up-regulation of the paralogous genes, CYFIP1 and ARPC1B. Expression of genes encoding subunits of the WAVE Arp2/3 complexes was overall not significantly altered in KO cell lines, nor was the expression of genes encoding Rac small GTPases (Fig.S6, Table S1).

Because of the differential phenotype of the triple KO between single cell and collective migration, we also measured a set of 9 EMT genes, *VIM*, *CDH1*, CDH2, *SNAI1*, *SNAI2*, *TWIST1*, *ZO1*, *ZEB1* and *ZEB2*. Among EMT-related genes, we found that the expression of vimentin and of the transcription regulator ZEB2 were strongly down-regulated in all KOs, including single KOs (Fig.4D, Fig.S6). ZEB2 controls expression of the vimentin gene in MCF10A cells (Bindels et al., 2006). Down-regulation of vimentin gene expression translated into strongly decreased protein levels in all KO clones, but this effect was not observed upon transient KD of CBA inhibitors (Fig.4E). This observation suggests that vimentin down-regulation was an adaptation to the absence of CBA inhibitors in the long term. When we examined vimentin expression and organization by immunofluorescence (Fig.4F), we found that vimentin was organized in a dense network in the majority of parental MCF10A cells. In contrast, single and double KOs had a decreased percentage of cells displaying such a network of vimentin filaments, and the triple KO even more so (Fig.4G). In wound healing assay, there was no absolute association between the leader cell phenotype and vimentin expression (Fig.4H). However, actively migrating leader cells in fingers were more likely to display a vimentin network than follower cells (Fig.4I).

### Vimentin opposes migration persistence of single cells

We wondered if vimentin down-regulation in KO cell lines was promoting or opposing the increased migration persistence of these lines. We first examined vimentin’s role in parental MCF10A using 2 independent siRNAs (Fig.5A). Vimentin depletion dramatically increased migration persistence of single cells (Fig.5B, Fig.S7, Movie S4), whereas overexpression of untagged vimentin cells did not yield any phenotype on persistence or other migration parameters of single cells (Fig.S8). In collective migration, vimentin-depleted MCF10A cells were less efficient at migrating than control cells, since they took longer to close the wound (Fig.5C). In the triple KO super-migrating clone, we isolated stable vimentin expressing cells to rescue their profound vimentin down-regulation (Fig.5D). As expected, these clones displayed a higher proportion of cells with vimentin networks than triple KO population (Fig.5E). Most importantly, migration persistence of these cells was efficiently rescued, albeit not completely (Fig.5F, Fig.S9). These results suggest that the transcriptional down-regulation of vimentin in super-migrating cells contributes to their exceptional migration persistence (Fig.6).

**Figure 5.**
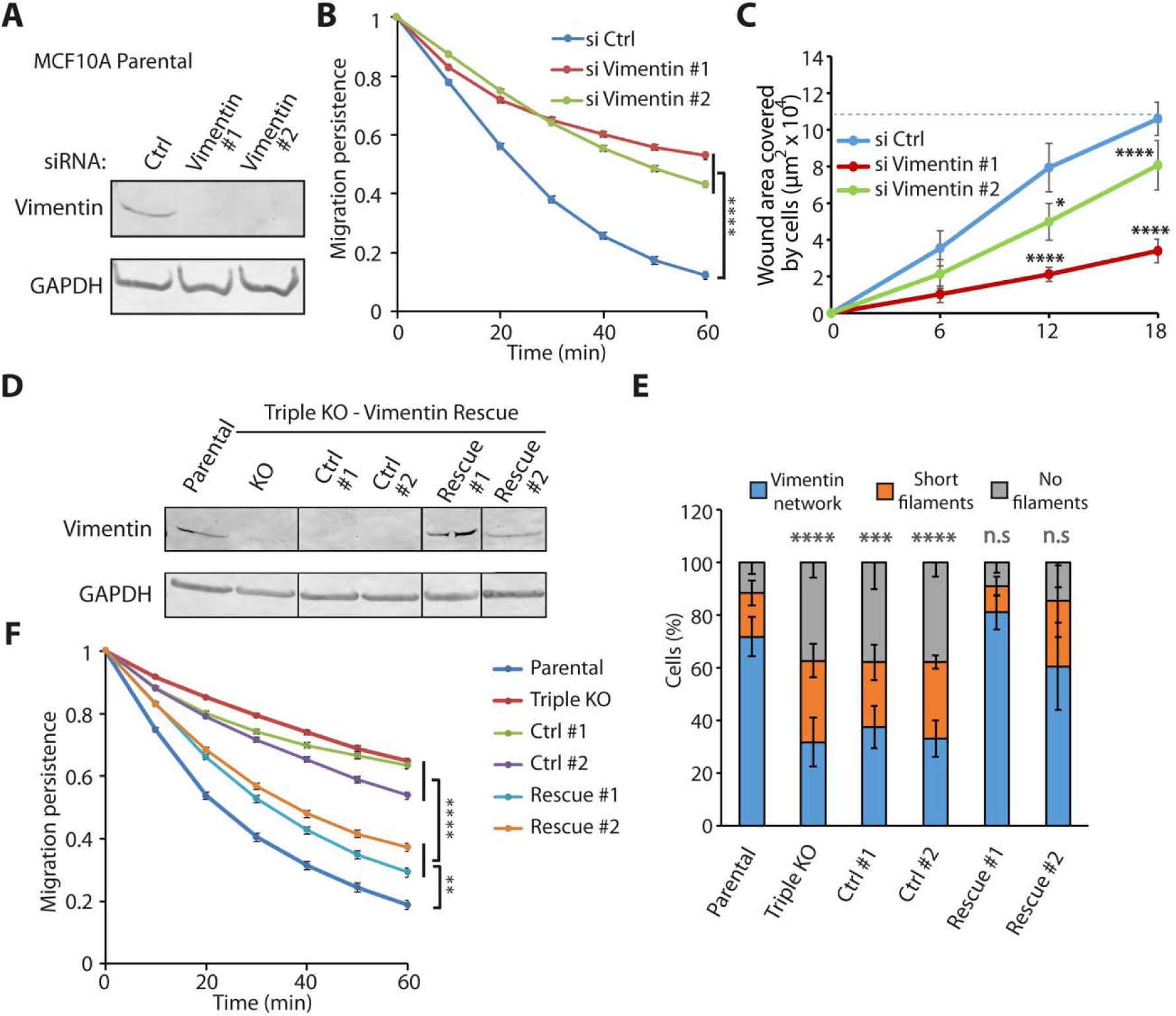
Vimentin antagonizes migration persistence of single cells. **A** MCF10A cells were depleted of vimentin using siRNAs. Western blots. **B** Migration persistence of single cells. 60 cells for each condition were pooled from 2 experiments. **C** Wound healing. The dashed line indicates full closure. 10 wounds for each condition were analyzed. 2 independent experiments gave similar results. **D** Western blots of parental, Triple KO and triple KO transfected by a plasmid expressing untagged vimentin or by an empty plasmid as a control. A single Western blot, where intervening lanes have been removed, is shown for each antibody. **E** Quantification of the percentage of cells displaying different types of vimentin organization. More than 400 cells per condition were taken into account. Kruskal-Wallis test. **F** Migration persistence of single cells. 65 cells for each condition were pooled from 2 experiments. ANOVA * p<0.05, ** p<0.01, *** p<0.001, **** p<0.0001, n.s. non-significant.

**Figure 6.**
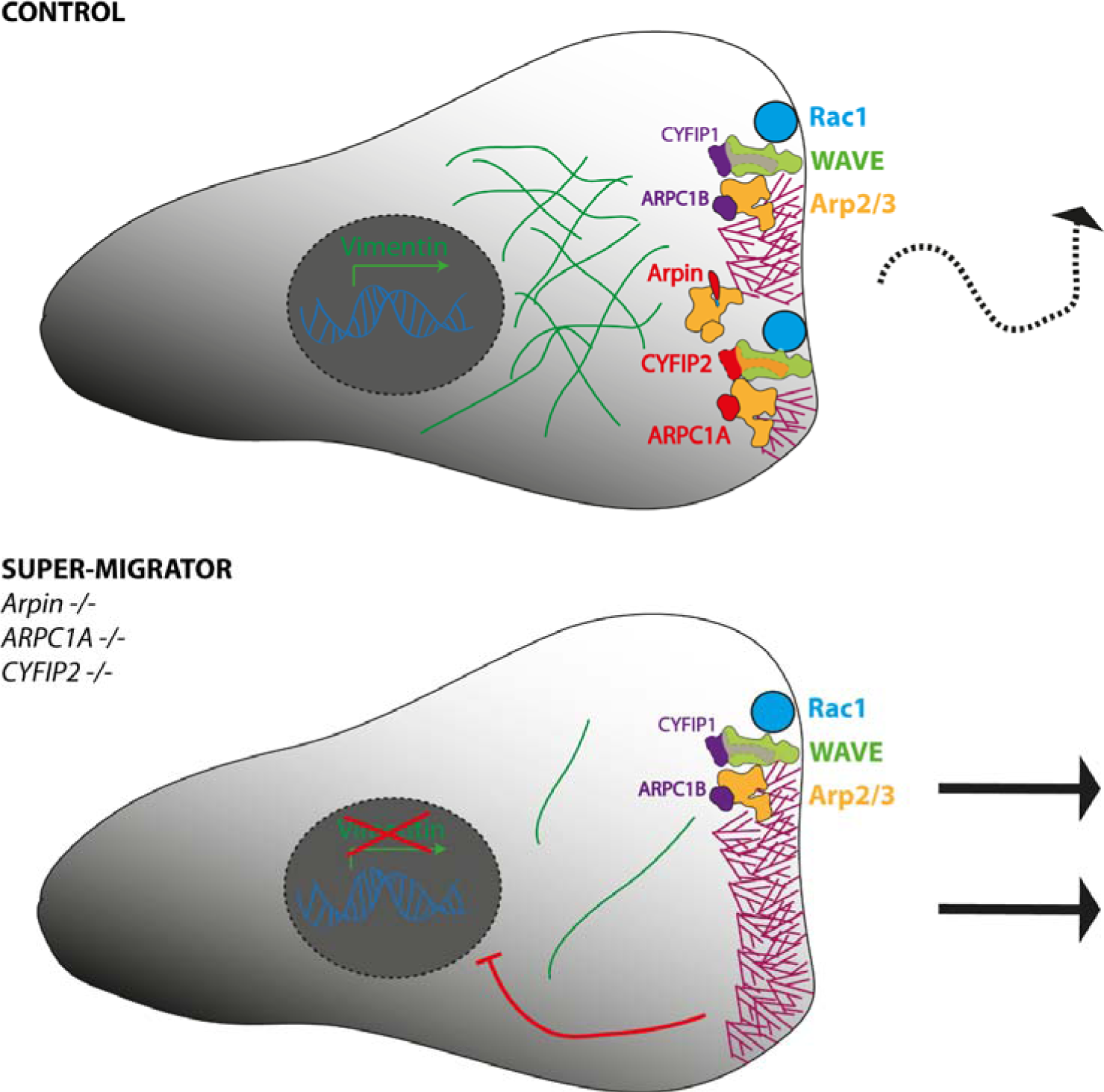
Model. In control cells, multiple molecular machines activated by Rac1 polymerizes branched actin in membrane protrusions. Active machines, CYFIP1-containing WAVE complexes and ARPC1B-containing Arp2/3 complexes, are balanced by relatively less active machines such as CYFIP2-containing WAVE complexes and ARPC1A-containing Arp2/3 complexes and by the Arp2/3 inhibitory protein Arpin. This balance results in moderately persistent cells. In super-migrator cells, where CYFIP2, ARPC1A and Arpin are inactivated, membrane protrusions are sustained by enhanced cortical branched actin and persistence of cell migration increases. Continuous overactivation of cortical branched actin in triple KO, but not in triple KD results in down-regulation of vimentin expression, which contributes to enhanced migration persistence.

## DISCUSSION

In this work, we combined the inactivation of 3 genes that antagonize migration persistence in MCF10A cells. In both KD and KO, there was no further effect when two genes were inactivated compared with a single inactivation, as if the system was buffered against a too dramatically enhanced persistence. Surprisingly enough, the addition of the third inactivation revealed a dramatically enhanced persistence in generated cells that we referred to as super-migrators, as if the third inactivation allowed to exceed a threshold. In zebrafish embryos, when we examined endodermal cells during gastrulation, the phenotype was roughly the same, since the triple inactivation generated cells that migrated more than any single or double KD. In the zebrafish system, however, an additional complexity was that the single ARPC1A KD did not generate a phenotype on its own, but amplified the persistence when combined with other KDs. Even if our super-migrators are very persistent, they are not yet similar to fish keratocytes, which are the most persistent vertebrate cells. To obtain the persistence and the characteristic morphology of fish keratocytes, it is perhaps required to combine our triple KO with inactivation of other negative regulatory genes. Among possible candidates, there are the Nance-Horan Syndrome protein NHSL1 (Law et al., 2021) and the Rac1 competitor CYRI (Fort et al., 2018).

The super-migrator phenotype of MCF10A cells was obtained with both triple KD and triple KO, but the mechanisms are different, because the long-term inactivation in KO results in gene expression changes. Using a targeted approach for 35 genes involved in the Rac-WAVE-Arp2/3 pathway or EMT, we identified down-regulation of ZEB2 and vimentin. This down-regulation was observed in any single KO indicating that it is a likely cell response to enhanced branched actin at the cortex. ZEB2 is a transcription factor the Zn finger family that was shown to control vimentin expression in MCF10A cells (Bindels et al., 2006). MCF10A cells are plastic epithelial cells, where cells in islets coexist with single cells, in regular culture conditions containing EGF. EGF induces vimentin expression in MCF10A cells (Bindels et al., 2006). We found that vimentin down-regulation was to a large extent responsible for the persistence of triple KO, even though this mechanism was not at play in the super-migrators obtained upon triple KD.

The down-regulation of ZEB2 and vimentin in super-migrators indicates that cells respond to enhanced cortical branched actin by becoming more epithelial and less mesenchymal. The role of vimentin in fibroblasts is very distinct from the role we uncover here in epithelial cells. Vimentin KO mouse embryonic fibroblasts were found to be less motile in vitro and in vivo upon wound healing (Eckes et al., 1998; Eckes et al., 2000). Unlike MCF10A cells, rat fibroblasts were less directionally persistent when vimentin was KO (Vakhrusheva et al., 2019). These opposite roles of vimentin in the persistence of fibroblasts and epithelial cells are striking. Along this line, it is interesting to note that the persistent fish keratocytes are also single cells dissociated from an epithelial monolayer (Rapanan et al., 2014), as if the transcriptional program of epithelial cells render cells more persistent than fibroblasts when these eptihelial cells happen to be isolated.

Gene expression changes induced by enhanced cortical branched actin are reminiscent of observations made with the exocyst subunit Exo70. Exo70 binds the Arp2/3 and contributes to its activation in the context of the Rac1-WAVE pathway and lamellipodium formation (Liu et al., 2012; Zuo et al., 2006). Exo70 is alternatively spliced and two splice forms, referred to as E for epithelial and M for mesenchymal, are differentially expressed in breast cancer cell lines as a function of their epithelial or mesenchymal morphology (Lu et al., 2013). Mesenchymal and invasive MDA-MB-231 cells express more of the M form of Exo 70, that more readily activates Arp2/3 than the E form. Switching expression to the E form of Exo70 induces ZEB2 expression in MDA-MB-231 cells and their mesenchymal-to-epithelial transition (Lu et al., 2013). Our work in MCF10A epithelial cells also showed that ZEB2 expression inversely correlates with cortical Arp2/3 activity and that enhancing cortical branched actin favors single cell migration over collective migration.

Super-migrator cells appeared less fit in collective behaviors than parental cells. Wound healing was less smooth with the apparition of leader cells at the front edge. Acinus morphogenesis was also affected. The triple KO cells differentiated into large acinus structures, as expected given the role of cortical branched actin in controlling cell cycle progression, but also into small acinus structures, which were abnormal in shape and extracellular matrix deposition. Importantly, despite their increased migration persistence as single cells, there was no sign of invasiveness, in line with the fact that these cells were driven towards a more epithelial program. Heterogeneity of the triple KO population was most evident at the level of vimentin expression and organization. Populations of cancer cells and aging cells are usually more heterogenous than populations of normal cells (Caspersson et al., 1963; Mahmoudi et al., 2019). Our work thus reveals that overactivation of the signaling pathway polymerizing cortical branched actin alters gene expression programs in a variable manner in the different cells of the population and renders epithelial cells less able to coordinate in collective behaviors.

## MATERIALS AND METHODS

### Cells and Transfections

The MCF10A cell line was cultured in DMEM/F12 medium (Gibco) supplemented with 5% horse serum (Sigma), 100 ng/mL cholera toxin (Sigma), 20 ng/mL epidermal growth factor (Sigma), 0.01 mg/mL insulin (Sigma), 500 ng/mL hydrocortisone (Sigma) and 100 U/mL penicillin/streptomycin (Gibco).

Human vimentin ORF (GenBank KU178388.1) was amplified from a vector provided by Dr. Alexander Minin (Institute of Protein Research of Russian Academy of Science). The PCR product was cloned into the home-made MXS CAG Blue SV40pA PGK Puro bGHpA between the Fsel and Ascl restriction sites and sequenced (Eurofins Genomics, Ebersberg, Germany) to ensure, that no mutations appeared after amplification. MCF10A cells were transfected using Lipofectamine 3000 (Invitrogen). Two days after transfection, 1 µg/mL Puromycin was added. Single clones were isolated by cloning rings, expanded and characterized.

For triple KD, cells were transfected using Lipofectamine RNAi max (Invitrogen) with 3 nM ARPC1A esiRNA (EHU105471, Sigma), 20 nM Arpin siRNA (5’-GUGGAUGUAUCUCGGCACA-3’, onTarget Plus, Dharmacon) and/or CYFIP2 (J-021477-05-0002, onTarget Plus, Dharmacon). Specific depletion was controlled with an equivalent amount of non-targeting siRNA (D-001810-01-05, onTarget Plus, Dharmacon) or GFP esiRNA (EHUEGFP, Sigma). siRNA-induced depletion of vimentin was obtained with 20 nM of siRNAs from Sigma, #1 5’-GUCUUGACCUUGAACGCAAdTdT-3’, #2 – 5’-GGUUGAUACCCACUCAAAAdTdT-3’ (Maier et al., 2015) and controlled with siCtrl–5’-AAUUCUCCGAACGUGUCACGUUU-3’ (Fokin et al., 2021). Cells were harvested or imaged after two or three days.

Knock-out cell lines were generated using the CRISPR/Cas9 system. The following gRNAs were used: negative control 5’-AAAUGUGAGAUCAGAGUAAU-3’; *ARPIN* 5’-GAGAACUGAUCGAUGUAUCU-3’; *ARPC1A* 5’-UAAGAAGAACGGGAGCCAGU-3’; *CYFIP2* 5’-CAUUUGUCACGGGCAUUGCA-3’. Cells were transfected with crRNA:tracrRNA duplex and the purified Cas9 protein by Lipofectamine CRISPRMAX™ Cas9 Transfection Reagent (all reagents were from Thermofisher Scientific). Cells then were subjected to dilution at 0.8 cells/well in 96 well plates. Clones were screened by Western blot. Targeted loci were amplified with the primers ARPINfor 5’-CCTGACAAGGTTCCTCCTGG-3’, ARPINrev 5’-TGCTGCTCAACACAGCCTTA-3’, ARPC1Afor 5’-ATTGACAGTTGTACGTGTCTCTG-3’, ARPC1Arev 5’-AAAGGAAGAGTGCCTGATTTGGA-3’, CYFIP2for 5’-GTTTCCACAGAGAGCTTGCG-3’, CYFIP2rev 5’-GGAGCTCAAGAAAGTGAGTAGTG-3’, and sequenced. In case of overlapping signals, PCR products were cloned (Zero Blunt PCR Cloning Kit, Thermofisher Scientific) and independent plasmids were sequenced to identify the 2 alleles.

### Individual migration of endodermal cells in zebrafish embryos

Embryos were obtained by natural spawning of Tg(−1.8gsc:GFP)ml1 adult fishes (Doitsidou et al., 2002). All animal studies were done in accordance with the guidelines issued by the Ministère de l’Education Nationale, de l’Enseignement Supérieur et de la Recherche and were approved by the Direction Départementale des Services Vétérinaires de l’Essonne and the Ethical Committee N°59.

Translation blocking morpholino (Gene Tool LLC Philomath) against ARPC1A (ATCTTCAAAGAATTTGCACCTCTGC) was designed for this study while morpholinos targeting CYFIP2 (CGACACAGGTTCACTCACAAAACAG) or Arpin (GTTGTCATAAATACGACTCATCTTC) were described earlier in Dang et al., 2013; Polesskaya et al., 2021.

Cells were forced to adopt an endodermal identity through the expression of the activated form of the Nodal receptor acvr1ba (acvr1ba*) (Peyriéras et al., 1998; Giger and David, 2017). Their nuclei were labeled through the expression of mCherry-tagged Histone2B. Corresponding mRNAs were synthesized using pCS2+ plasmids containing acvr1ba* or Histone2B-mCherry sequence and the mMessage mMachine SP6 kit (Thermo Fischer).

Donor embryos were injected at the 8-cell stage with 0.2 nl of a solution containing acvr1ba* (0.6 ng/µl), Histone2B-mCherry mRNA (30 ng/μl) and morpholinos against ARPC1A (0.2 mM), Arpin (0.2 mM) or CYFIP2 (0.2 mM), alone or in combination. Small groups of GFP expressing endodermal cells were transplanted at the shield stage (6 hpf) to the animal pole of an untreated host (Giger and David, 2017; Boutillon et al., 2018). Embryos were then cultured in embryo medium with 10 U/mL penicillin and 10 μg/mL streptomycin. Transplanted embryos were mounted in 0.2% agarose in embryo medium and imaged between shield stage and 85% epiboly (6 - 9 hpf) under an inverted TCS SP8 confocal microscope (Leica) equipped with environmental chamber (Life Imaging Services) at 28°C, using a HCX PL Fluotar 10x/0.3 objective (Leica). Visualization of 3D movies and nuclei tracking were done using Imaris (Bitplane). Cell migration parameters were extracted using custom codes in Matlab (Math Works) (Boutillon et al., 2022) and autocorrelation was computed using published Excel macros (Gorelik and Gautreau, 2014).

### Antibodies

For Western blots, the following antibodies were used: CYFIP2 (Sigma SAB2701081), Tubulin clone DM1A (Sigma T9026), ARPC1A (Sigma #HPA004334), Arpin home-made (Dang, Nature), p62/DCTN4 clone H-4 (Santa Cruz Biotechnology). For immunofluorescence anti-Laminin V, clone D4B5 (Merck). For both applications, Anti-Vimentin, clone V9 (sc-6260, Santa Cruz) was used. Secondary goat anti-mouse and anti-rabbit antibodies conjugated with Alexa Fluor 488, 555 and 647 used for immunofluorescence were from Life Technologies. Secondary goat anti-mouse and anti-rabbit antibodies conjugated with alkaline phosphatase used for Western blots were from Promega.

### Western Blots

Cells were lysed in 50 mM Hepes, pH7.7, 150 mM NaCl, 1 mM CaCl2, 1% NP40, 0.5% Na Deoxycholate, and 0.1% SDS supplemented with a protease inhibitor cocktail (Roche). Lysates were spun at +4℃ and 20000 xg and supernatants were mixed with LDS (ThermoFischer) and 2.5% of β-ME. SDS-PAGE was performed using NuPAGE 4–12% Bis-Tris gels (ThermoFischer Scientific). After transfer, nitrocellulose membranes were blocked in 5% skimmed milk incubated with primary and secondary antibodies conjugated with alkaline phosphatase and developed with NBT/BCIP as substrates (Promega).

### qRT-PCR

Total RNA from cell lines was extracted by NucleoSpin RNA Plus Kit (Macherey-Nagel). Specific mRNAs were quantified from the cycle number (Ct value) at which the increase in the fluorescence signal started to be detected by the laser detector of the QuantStudio 7 Flex Real-Time PCR System (Thermo Fisher Scientific, Waltham, MA, USA) as previously described (Bieche et al., 2001). Gene expression levels were normalized to *TBP* expression levels (NM_003194) used as an endogenous RNA control and to the control condition, MCF10A parental cell line, to calculate the fold change. Nucleotide sequences of the primers used are detailed in Table S2.

### Videomicroscopy of Cell Migration

Videos of cell migration were acquired on an inverted Axio Observer microscope (Zeiss) equipped with a Pecon Zeiss incubator XL multi S1 RED LS (Heating Unit XL S, Temp module, CO2 module, Heating Insert PS and CO2 cover), a definite focus module and an ORCA-Flash4.0 V3 Digital CMOS camera. Pictures were taken every 5 or 10 min for 24 h using the Plan-Apochromat 20X/0.80 or Plan-Apochromat10X/0.40 air objectives. For wound healing, wounds were produced in a cell monolayer either by a pipet tip or by removing inserts from μ-dishes 35 mm (Ibidi). Cells were imaged every 20 minutes during 24 hours.

### Acinus formation and staining

MCF10A cells were seeded on top of polymerized matrigel (CB-40230C, Corning) in Millicell EZ SLIDE 8-well glass chamber slide (PEZGS0816, Millipore) in a medium containing 4ng/mL EGF (4ng/mL) and 1% serum and supplemented with 2% of matrigel. During 3 weeks, medium was regularly changed. Acini were then fixed in 2% PFA in PBS permeabilized with 0.5% Triton X-100, rinsed with PBS/glycine (130mM NaCl, 7mM Na2HPO4, 3.5mM NaH2PO4, 100mM glycine), blocked in IF Buffer (130mM NaCl, 7mM Na2HPO4, 3.5mM NaH2PO4, 0.1% bovine serum albumin, 0.2% Triton X-100, 0.05% Tween-20) + 10% FBS first and then with IF Buffer + 10% FBS + 20 µg/ml goat anti-Rabblit Fc fragment (111-005-046, Jackson ImmunoResearch). Acini were incubated with the primary antibodies in the secondary block solution washed with IF buffer and then incubated with secondary antibody in IF Buffer +10% FBS. Then acini were incubated with DAPI, rinsed with IF buffer and Mounted with Abberior Mount Liquid Antifade (Abberior) and sealed with nail polish. Overview images of acini were taken on an Olympus CKX53 microscope, equipped with a DP22 camera (Olympus) and DP2-SAL firmware (Olympus). Acinus size was then calculated in Fiji by manually contouring acini.

### Immunofluorescence

Cells were seeded on glass coverslips coated with 20 µg/mL bovine fibronectin (Sigma) for 1 h at 37 °C. Cells were fixed in PBS/3.2% PFA then quenched with 50 mM NH_4_Cl, permeabilised with 0.5 % Triton X-100, blocked in 2% BSA and incubated with antibodies (1-5 µg/mL for the primary, 5 µg/mL for the secondary) or 1:3000 diluted SiR-actin (Tebu-bio). Nuclei were counter stained with DAPI (Live Technologies). Images were acquired by Axio Observer microscope (Zeiss). Images of acini were obtained on a SP8ST-WS confocal microscope equipped with a HC PL APO 63x/1.40 oil immersion objective, a white light laser, HyD and PMT detectors.

### Migration Analysis

Image analysis was performed in ImageJ or FIJI software. In the single cell migration assay, only cells that were freely migrating for 7 h or more were taken into account. Cell trajectories were acquired with the ImageJ Manual tracking plugin. Random migration of single cells and migration persistence, based on the angular shift between frames, was analyzed as previously described (Dang et al., 2013) using the DiPer program (Gorelik and Gautreau, 2014). To assess speed of wound closure, wound areas were manually drawn and measured at different time points. Variance of migration (St-Dev/Mean) in wound healing was calculated by measuring the distance moved by the leading edge perpendicular to the initial wound edge, for each pixel along the wound. To this end, time-lapse images were first thresholded, segmented in FIJI and analyzed using custom-made MATLAB scripts.

### Statistics

The autocorrelation curves corresponding to migration persistence were analyzed in R and fitted by an exponential decay as described (Boutillon et al., 2022)

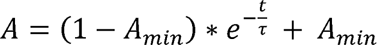

where A is the autocorrelation, *t* the time interval, A_min_ the plateau, and τ the time constant of decay. The plateau value A_min_ is set to zero for the cell lines in vitro, as they do not display overall directional movement. Time constants τ, reflecting directional persistence, were then compared using one-way ANOVA on non-linear mixed-effect models to take into account the resampling of the same statistical unit.

Other statistical analyses were carried out with GraphPad Prism software (v7.00) and Microsoft Excel 2016. When not stated otherwise, ANOVA and Kruskal-Wallis test were used. Shapiro-Wilk normality test was applied to examine whether data fit a normal distribution. If data satisfied the normality criterion, ANOVA was followed by a post-hoc Tukey’s multiple comparison test. If not, a non-parametric Kruskal-Wallis test was followed by a post-hoc Dunn’s multiple comparison test. 4 levels of significance were distinguished: * p<0.05, ** p<0.01, *** p<0.001, **** p<0.0001.

## Abbreviations

CBA: Cortical Branched Actin
KO: Knock-Out
KD: Knock-Down
indel: insertion/deletion
ORF: Open Reading Frame
MSD: Mean Square Displacement
EMT: epithelial-mesenchymal transition
NPF: Nucleation Promoting Factor
SD: Standard Deviation
SEM: Standard-Error-to-Mean.

## Author Contributions

AIF performed most experiments with human cells. JJ and YW contributed to migration analyses. GS and AP contributed to KO clone isolation and performed statistical analyses of migration persistence. AB and NBD performed cell migration experiments in Zebrafish. LC, SV and IB performed qRT-PCR analyses. NBD, IB and AMG performed supervision in their respective groups. AIF and AMG conceived the study, coordinated the work and wrote the manuscript.

## Acknowledgements

We thank Dmitry Guschin and Pierre Mahou for excellent technical assistance, Alexander Minin for providing vimentin cDNA. This work was supported by grants from Agence Nationale de la Recherche (ANR-20-CE13-0016 to AMG and NBD and ANR-22-CE13-0041 to AMG), Fondation ARC pour la Recherche sur le Cancer (ARC PJA 2021 060003815 to AMG), from Institut National du Cancer (INCA_11508 for AMG and IB). We thank the Polytechnique Bioimaging Facility for confocal microscopy partly supported by Région Ile-de-France (interDIM) and Agence Nationale de la Recherche (ANR-11-EQPX-0029 Morphoscope2, ANR-10-INBS-04 France BioImaging).

